# Whole-Genome Enrichment and Sequencing of *Chlamydia trachomatis* Directly from Patient Clinical Vaginal and Rectal Swabs

**DOI:** 10.1101/2020.09.04.282459

**Authors:** Katherine E. Bowden, Sandeep J. Joseph, John Cartee, Noa Ziklo, Damien Danavall, Brian H. Raphael, Timothy D. Read, Deborah Dean

## Abstract

*Chlamydia trachomatis* is the most prevalent cause of bacterial sexually transmitted infections (STIs) worldwide. U.S. cases have been steadily increasing for more than a decade in both the urogenital tract and rectum. *C. trachomatis* is an obligate intracellular bacterium that is not easily cultured, limiting the capacity for genome studies to understand strain diversity and emergence among various patient populations globally. While Agilent SureSelect^XT^ target-enrichment RNA bait libraries have been developed for whole-genome enrichment and sequencing of *C. trachomatis* directly from clinical urine, vaginal, conjunctival and rectal samples, efficiencies are only 60-80% for ≥95-100% genome coverage. We therefore re-designed and expanded the RNA bait library to augment enrichment of the organism from clinical samples to improve efficiency. We describe the expanded library, the limit of detection for *C. trachomatis* genome copy input, and the 100% efficiency and high-resolution of generated genomes where genomic recombination among paired vaginal and rectal specimens from four patients was identified. This workflow provides a robust approach for discerning genomic diversity and advancing our understanding of the molecular epidemiology of contemporary *C. trachomatis* STIs across sample types, among geographic populations, sexual networks, and outbreaks associated with proctitis/proctocolitis among women and men who have sex with men.

**Importance:** *Chlamydia trachomatis* is an obligate intracellular bacterium that is not easily cultured, and there is limited information on rectal *C. trachomatis* transmission and its impact on morbidity. To improve efficiency of previous studies involving whole genome target enrichment and sequencing of *C. trachomatis* directly from clinical urine, vaginal, conjunctival, and rectal specimens, we expanded the RNA bait library to augment enrichment of the organism from clinical samples. We demonstrate an increased efficiency in the percentage of reads mapping to *C. trachomatis*. We show the new system is sensitive for near identical genomes of *C. trachomatis* from two body sites in four women. Further, we provide a robust genomic epidemiologic approach to advance our understanding of *C. trachomatis* strains causing ocular, urogenital and rectal infections, and to explore geo-sexual networks, outbreaks of colorectal infections among women and men who have sex with men, and the role of these strains in morbidity.

## Introduction

*Chlamydia trachomatis* is the most common cause of bacterial sexually transmitted infections (STIs) worldwide and the most notifiable disease in the United States^1,2^. Although *C. trachomatis* infection can present with conjunctivitis, pharyngitis, urethritis, vaginal discharge, proctitis or inguinal syndrome, most infections are asymptomatic, which can lead to reproductive morbidity in women and proctitis or proctocolitis in women and men who go untreated^2^.

*C. trachomatis* strains are classified by genotype based on the outer membrane protein gene (*omp*A), which encodes the major outer membrane protein (MOMP), and is typically linked to clinical presentation^3^. Twenty one genotypes are grouped according to disease association: ocular (A-C, Ba); urogenital and anorectal (D-K, Da, Ia, Ja); and lymphogranuloma venereum (LGV; L_1_-L_3_, L_2_a, L_2_b, L_2_c)^2,4–7^. Several studies have reported that the prevalence of these genotypes differ by anatomical site and sexual network. Genotypes D and G are more commonly detected in the anorectal tract and, along with genotype J, are prevalent in women and men who have sex with men (MSM)^8–15^. Genotypes D, E, and F are found in the majority of urogenital infections and are common among heterosexuals^8,16^. LGV genotypes are prevalent in HIV infected MSM and associated with an anorectal infection and the inguinal syndrome^17–21^.

A 2016 meta-analysis of extragenital *C. trachomatis* and *Neisseria gonorrhoeae* infections in women, MSM, and men who have sex only with women (MSW) demonstrated a median prevalence of 8.7%, 8.9%, and 7.7%, respectively, for rectal *C. trachomatis*, with infection often being asymptomatic^22^. Further, a number of studies have shown that rectal infections outnumber those in the urogenital tract of women and are on the increase among MSM^23–28^. Although common, rectal *C. trachomatis* transmission and its impact on morbidity is not well understood, likely due to the lack of routine screening of populations other than MSM^1,22,29^. In May of 2019, the FDA cleared two diagnostic tests, the Aptima Combo 2 Assay (Hologic, Inc.) and the Xpert CT/NG (Cepheid), for the use of extragenital specimens in the detection of *C. trachomatis* and *N. gonorrhoeae*. This recent diagnostic advancement will improve screening and surveillance capacity while offering an opportunity to better understand transmission of rectal *C. trachomatis* and its role in morbidity.

Transmission and molecular epidemiologic studies of *C. trachomatis* rely on *ompA* genotyping, multi-locus sequence typing (MLST), and multi-locus variable number tandem repeat analysis (MLVA)^10,30–39^. Unfortunately, these techniques are challenging and laborious when performed on clinical specimens and, except for *ompA* genotyping, often require tissue culture to generate sufficient DNA. Further, due to low genetic resolution, these methods fail to demonstrate precise inter- and intra-strain recombination events across the genome that contribute to strain diversity^40–44^. Recombination has been important in creating emerging strains of *C. trachomatis* such as L_2_b among MSM in many countries of the world and recombinant strains L_2_/D (termed L_2_c) and L_2_b/D-Da in the U.S. and Portugal, respectively^28,40–47^.

Enrichment of the low copy number of *C. trachomatis* in clinical specimens presents the greatest challenge for culture-independent genome sequencing^48^. Initially, this methodology employed immunomagnetic separation (IMS) for enrichment, followed by genome amplification using multiple displacement amplification (MDA), but the demonstrated success rate (15-30%) was low across clinical specimens^49–51^. Other methodologies currently in use include depletion-enrichment, cell-sorting-MDA, and multiplexed microdroplet PCR^48^. In 2014, Christiansen et al. sequenced *C. trachomatis* from urine and vaginal swabs with an 80% (8/10) success rate (≥95-100% coverage of the respective reference genome) using custom RNA baits to enrich for *C. trachomatis* during library preparation^52^. The same RNA bait library was subsequently used for conjunctival samples with a 60% (12/20) success rate and similar genome coverage. More recently, another Agilent custom RNA bait library was developed for *C. trachomatis* enrichment from rectal samples^53^. This RNA bait methodology has therefore been used to understand genomic diversity in circulating *C. trachomatis* ocular, urogenital and LGV lineages from clinical specimens but with varying success^53–57^.

In an effort to make direct sequencing of all *C. trachomatis* strains causing clinical infections more efficient, we designed an expanded RNA bait library (based on 85 *C. trachomatis* complete genomes, with 34,795, 120-mer probes instead of 74 genomes with 33,619, 120-mer probes as for previous bait libraries^48,52^) and optimized the experimental protocol. The new system will increase our ability to sequence *C. trachomatis* genomes of current circulating strains causing ocular, urogenital and rectal infections in diverse populations and will provide a more robust data set to understand current sexual networks and transmission within and among anatomic sites. From these data, we will be able to differentiate clusters and sporadic cases during outbreaks, and potentially identify novel markers for typing *C. trachomatis* in addition to exploring the role of these strains in ocular, genital and colorectal morbidity.

## Methods

### Spiked serial dilutions of *C. trachomatis* reference strain D/UW-3/Cx

Genomic DNA (gDNA) from *C. trachomatis* reference strain genotype D/UW-3/Cx (ATCC # VR-885D) was used to determine the limit of detection (LOD) for this workflow. The LOD was set at the minimal genome copy number required to generate a ≥5X read depth with ≥98% genome coverage compared to the reference strain of the same *ompA* genotype. Six 100 μL serial dilutions (10^−1^ to 10^−6^) were prepared by spiking into 1x PBS. A standard curve based on ATCC’s reported copy number for genotype L_2_b (ATTC # VR-902BD) was generated using the real-time polymerase chain reaction (RT-PCR) targeting the *C. trachomatis* single-copy polymorphic membrane protein gene *pmpH*^58^. This standard curve (y = 1 × 10^e(−0.602x)^; R^2^ = 0.987) was then used to calculate a more precise genome copy number for each serial dilution of genotype D.

### Spiked mock samples of *C. trachomatis* genotype L_2_b

Genomic DNA (gDNA) from *C. trachomatis* clinical strain genotype L_2_b (ATCC # VR-902BD) was used to ensure success of this workflow with a *C. trachomatis* strain prevalent in anorectal infections in HIV infected MSM. Three 100 μL serial dilutions (9,000-900,000 total genome copies) were prepared by spiking into 1x PBS, all within the LOD established using *C. trachomatis* reference strain genotype D.

### *C. trachomatis* clinical specimens and determination of *C. trachomatis* genome copy number

Clinical urogenital and rectal samples were obtained from women aged 18 to 40 years who were at high risk for STIs after informed consent as part of a separate study that was approved by UCSF Benioff Children’s Hospital Oakland Research Institute IRB. For this study, the samples were deidentified of all personal identifiers with no trace to the patient names. FLOQswab vaginal and rectal swabs (Copan, Murietta, CA) had been collected using standard techniques by trained clinic staff and screened for *C. trachomatis* using the Xpert CT/NG test (Cepheid, Sunnyvale, CA). Four clinical vaginal samples and the four paired rectal swabs from the same four women (randomly selected using a table of random numbers from over 200 women positive for *C. trachomatis* at both sites) were used in this study.

Approximately 200 μL of remnant swab collection buffer that had not been run in the Xpert test was lysed with 59 μL of a cocktail consisting of 50 μL lysozyme (10 mg/mL; MilliporeSigma, St. Louis, MO), 3 μl of lysostaphin (4,000 U/mL in sodium acetate; MilliporeSigma) and 6 μl of mutanolysin (25,000 U/mL; MilliporeSigma) for 1 hour at 37°C as described^59^. gDNA was then purified from the lysate using the QIAamp DNA mini kit (Qiagen) according manufacturer’s instructions. For rectal swabs collected in M4 media (Thermo Fisher, South San Francisco, CA), 200 μL was treated as above.

gDNA was subjected to an in-house qPCR to quantitate the genomic copy number of *C. trachomatis* as we have described^60^. Briefly, a standard curve was calculated based on 10-fold serial dilutions of a linearized plasmid containing the single copy *ompA* gene. *C. trachomatis* genomic copy number of the clinical samples was determined based on the standard curve.

### *C. trachomatis ompA* genotyping and plasmid sequencing

*ompA* genotyping was performed as previously described^61^. Briefly, primers flanking the *ompA* gene were used for PCR. The PCR product was purified by exoSAP-IT (Thermo Fisher) and subjected to Sanger sequencing using the PCR primers^6161^. Forward and reverse sequences were aligned using MAFFT v7.450^62^ to create a consensus sequence that was aligned to all reference *ompA* genotypes. The reference strains included A/HAR-13, B/TW-5/OT, Ba/Apache-2, C/TW-3/OT, D/UW-3/Cx, Da/TW-448, E/Bour, F/IC-Cal-13, G/UW-57/Cx, H/UW-4/Cx, I/UW-12/Ur, Ia/UW-202, J/UW-36/Cx, Ja/UW-92, K/UW-31/Cx, L_1_/440, L_2_/434, L_2_a/UW-396, L_2_b/UCH-1/proctitis, L_2_c, and L_3_/404.

Five overlapping PCR primer pairs were designed using the IDT PrimerQuest® Tool (https://www.idtdna.com/pages/tools/primerquest) to amplify the entire plasmid (Supplementary Table ST1). The thermocycling parameters were: 3 min at 95°C followed by 40 cycles of 95°C for 30 sec, 56°C for 30 sec and 72°C for 1 min 10 sec with a final incubation at 72°C for 7 min. The PCR product was purified and sequenced as above, and the consensus sequence was aligned to all reference plasmid sequences as above using MAFFT v7.450.

### Quantification and fragmentation

Samples were quantified using the Qubit® 2.0 Fluorometer, and human gDNA (Promega, San Luis Obispo, CA) was added to reach a total input of 3 μg/130uL for fragmentation and library prep. Samples were sheared on the Covaris® LE220plus using the 8 microTUBE strip V1 (PM# 520053; Covaris, Woburn, MA) with the base pair (bp) mode set to 250-300 bp following the manufacturer’s instructions.

### RNA Bait library design

A 2.698 Mbp RNA bait library consisting of 34,795 120-mer probes spanning 85 GenBank *C. trachomatis* reference genomes were designed using Agilent SureDesign. No plasmid probes were included in the RNA bait library construction. The bait library was synthesized by Agilent Technologies. The custom designed RNA bait library (ELID: 3173001) used in this study can be retrieved by contacting Agilent Technologies, INC (Santa Clara, CA).

### Library prep

After shearing, the SureSelect^XT^ Target Enrichment System for Illumina Paired-End Multiplexed Sequencing Library (VC2 Dec 2018) and all recommended quality control steps were performed on all gDNA samples. A 16-hour incubation at 65°C was performed for RNA bait library hybridization. Post-capture PCR cycling was set at 12 cycles based on a capture library size > 1.5 Mb.

### Illumina MiSeq sequencing

The eight clinical samples were multiplexed for two runs of paired end sequencing on an Illumina MiSeq using a 300 bp v2 reagent kit. For the final multiplexed library pool, libraries were diluted to 2 nM/3 μL in Low TE for a final concentration of 10 pM; 12.5 pM PhiX was added to the final pool that was loaded onto the MiSeq. All sequencing data associated with this study was submitted to the National Center for Biotechnology Information’s sequence read archive (SRA) under the BioProject accession ID: PRJNA609714.

### Sequence and phylogenetics analysis

Host genome sequences were first filtered from the raw sequencing dataset using Bowtie2 version 2.2.9^63^, which removed any contaminating human sequences using the h19 human reference genome^64^. Cutadapt version 1.8.3^65^ was used to trim specified primers and adaptors, and to filter out reads below Phred quality scores of 15 and read length below 50 bps. Deduplication of the reads were performed using Clumpify (sourceforge.net/projects/bbmap/) with dedupe=t option to prevent biased coverage of genomic regions. *C. trachomatis* sequencing reads were selected using K-SLAM^66^, a k-mer – based metagenomics taxonomic profiler, which used a database containing all bacterial and archaeal reference nucleotide sequences. The presence of *C. trachomatis* sequences was also confirmed using Metaphlan2^67^. We generated a custom version of the *C. trachomatis* D/UW-3/CX reference sequence (NC_000117.1) from which we masked 6 rRNA genes (CT_r01 (16SrRNA_1), CT_r02 (23SrRNA_1), CT_r03 (5SrRNA_1), CT_r04 (16SrRNA_2), CT_r05 (23SrRNA_2) and CT_r06 (5SrRNA_2) present in the repeated rRNA operons using bedtools v2.17.0^68^ ‘maskfasta’. Prefiltered chlamydial reads were mapped against this custom reference genome using BWA mem v2.12.0^69^ (MapQ ≥ 20), followed by consensus sequence generation and estimation of sequencing depth and mapping statistics using samtools^70^ (options ‘depth’ and ‘mpileup’) and bcftools v1.9. The prefiltered *C. trachomatis* sequencing reads were also used to generate *de novo* short read assemblies using SPAdes 3.7.0^71^ with the “careful” option. To genotype the patient samples, *de novo* contigs were used to extract and compare the *ompA* genes against a customized BLAST^72^ database of the 21 reference *ompA* sequences (see above).

For phylogenetic analysis, apart from the patient genome sequences (n=8), we also included all *C. trachomatis* genomes (without plasmid sequences) available in NCBI (n=94) and used a reference mapping approach with the above-mentioned custom version of the *C. trachomatis* D/UW-3/CX reference genome sequence. In short, full length whole genome alignments were generated using Snippy v3.1 (https://github.com/tseemann/snippy), which identifies variants using Freebayes v1.0.2^73^ with a minimum 10X read coverage and 90% read concordance at a locus for each single nucleotide polymorphism (SNP). Regions of increased density of homoplasious SNPs introduced by possible recombination events were predicted iteratively and masked using Gubbins^74^. Final phylogenetic tree was reconstructed using RAxML^75^ on the recombination removed alignment using the General Time Reversible (GTR) model. The genes located within the putative recombination blocks for the patient samples were identified by comparing the alignment genomic co-ordinates for the predicted recombination blocks to the gene annotations of the reference genome. Within host SNP differences were derived from the alignment before masking the predicted recombination events.

## Results

### Limit of detection

The limit of detection (LOD) was determined using varying genomic copy numbers of a *C. trachomatis ompA* genotype D strain mapped to the D reference strain. Genomic libraries were prepared, enriched for *C. trachomatis* and sequenced from spiked serial dilutions of gDNA bulked with human DNA for a total input of 3 μg DNA (for fragmentation and library preparation) using the expanded RNA bait library and Agilent SureSelect^XT^ protocol. Total *C. trachomatis* genome copy input ranged from 265-7,854,406 copies, which is a similar range to genome copies from clinical samples (see below), with 1.33-99.28% of the quality-controlled reads binned as *Chlamydia* species, along with a mean mapping read depth to *C. trachomatis* reference genome ranging from 0.57-562.67, respectively (Table 1, Figure 1, Supplementary Figure S1). With the QC criteria for efficiency set at ≥ 98% genome coverage at ≥ 5X read depth, genotype D had an LOD of 16,945 total genome copies (Table 1, Figure 1).

**Table 1:**
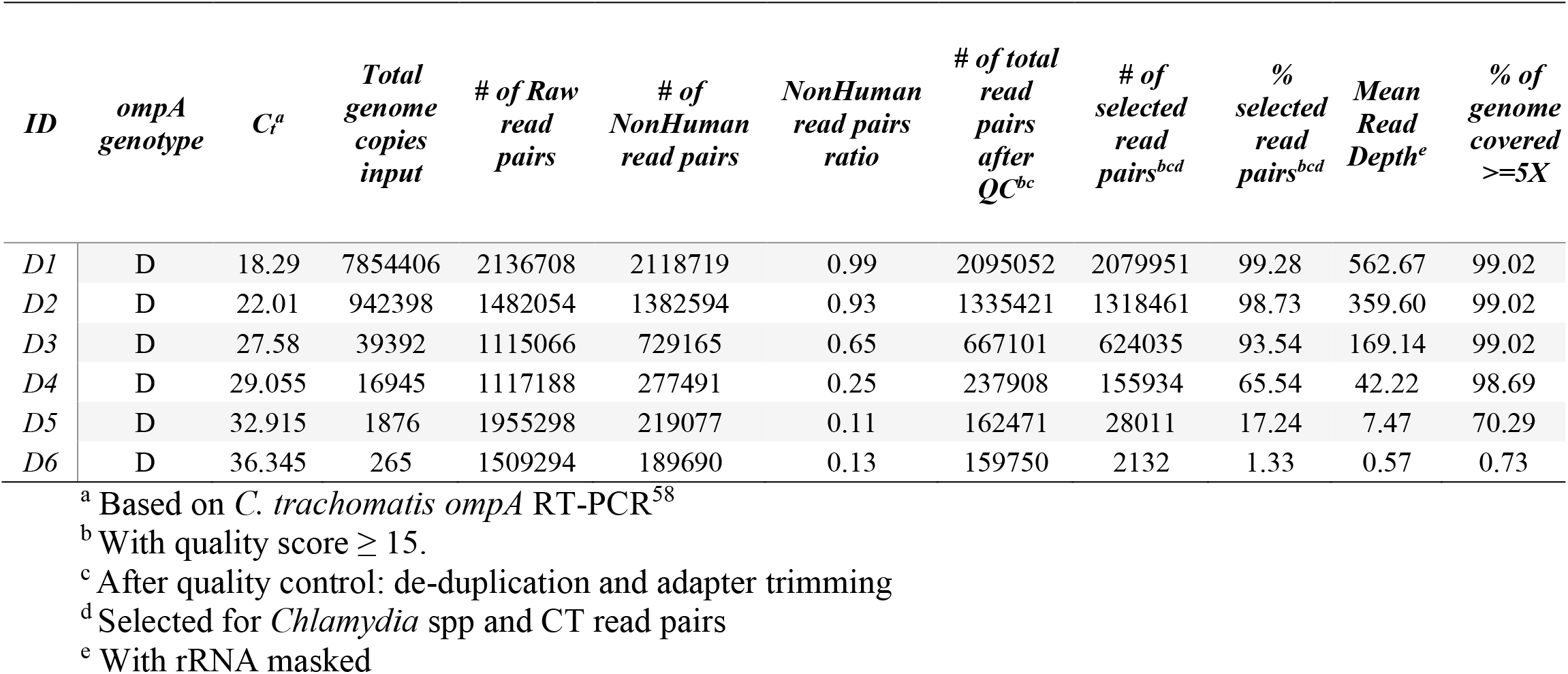
Sequence data analysis of *C. trachomatis* gDNA from spiked serial dilutions of reference strain D/UW-3/Cx.

**Figure 1:**
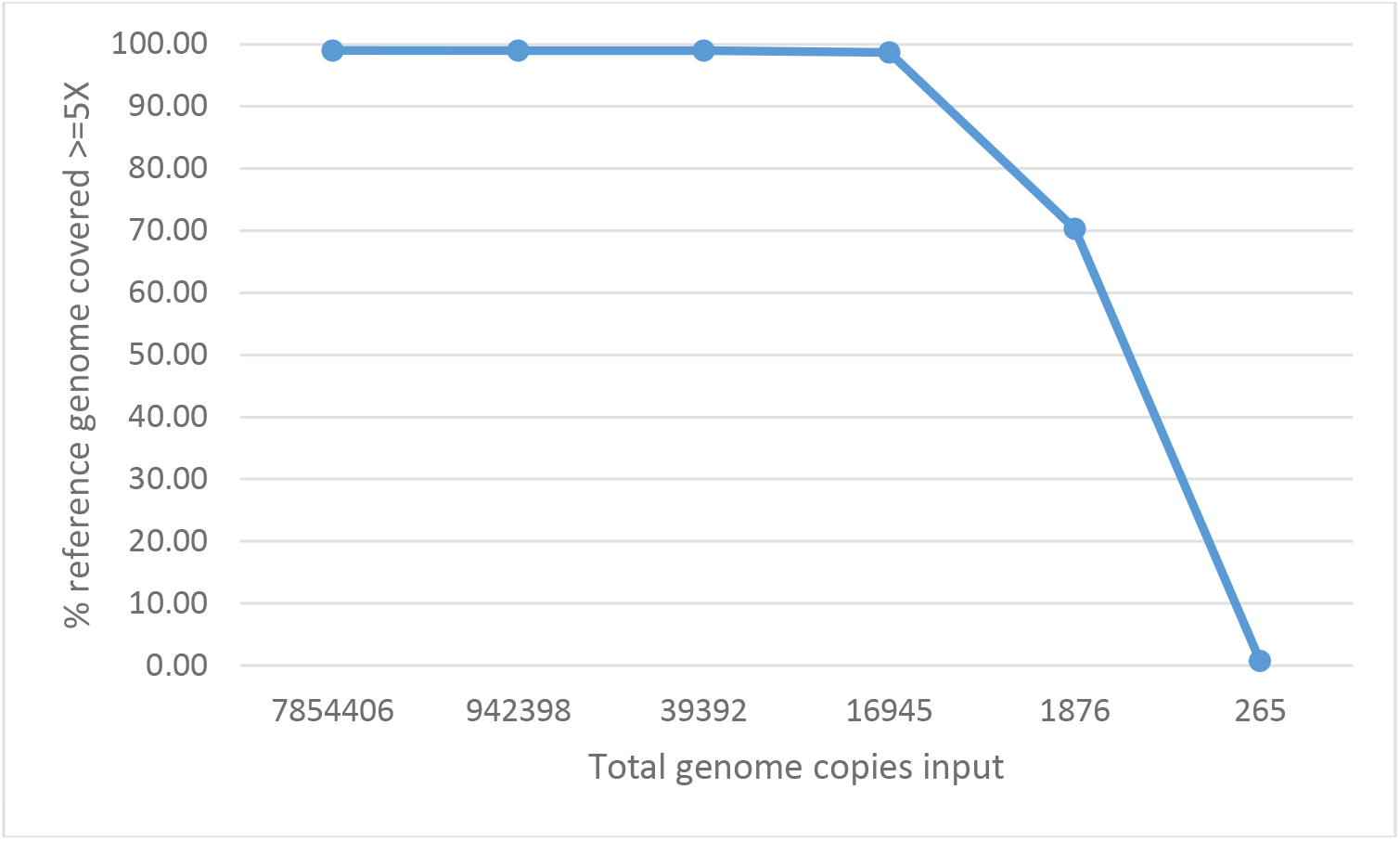
Limit of detection (LOD) of genome copies using the SureSelect^XT^ target-enrichment workflow for spiked serial dilutions of reference strain D/UW-3/Cx genomic DNA (gDNA). The percent coverage of the reference genome for each serial dilution is represented as blue line.

### Enrichment and genomic sequencing of *C. trachomatis* from genotype L_2_b and patient specimen sets

To ensure efficiency of target-enrichment from a *C. trachomatis ompA* genotype prevalent in anorectal infections in HIV infected MSM, genomic libraries from spiked mock samples of genotype L_2_b were prepared, enriched for *C. trachomatis* and sequenced. Total *C. trachomatis* genome copy input ranged from 9,000-900,000 copies with 97.22-98.77% of the quality-controlled reads binned as *Chlamydia* species along with a mean mapping read depth to *C. trachomatis* reference genome ranging from 531.81-550.78, respectively (Table 2, Supplementary Figure S2).

**Table 2:**
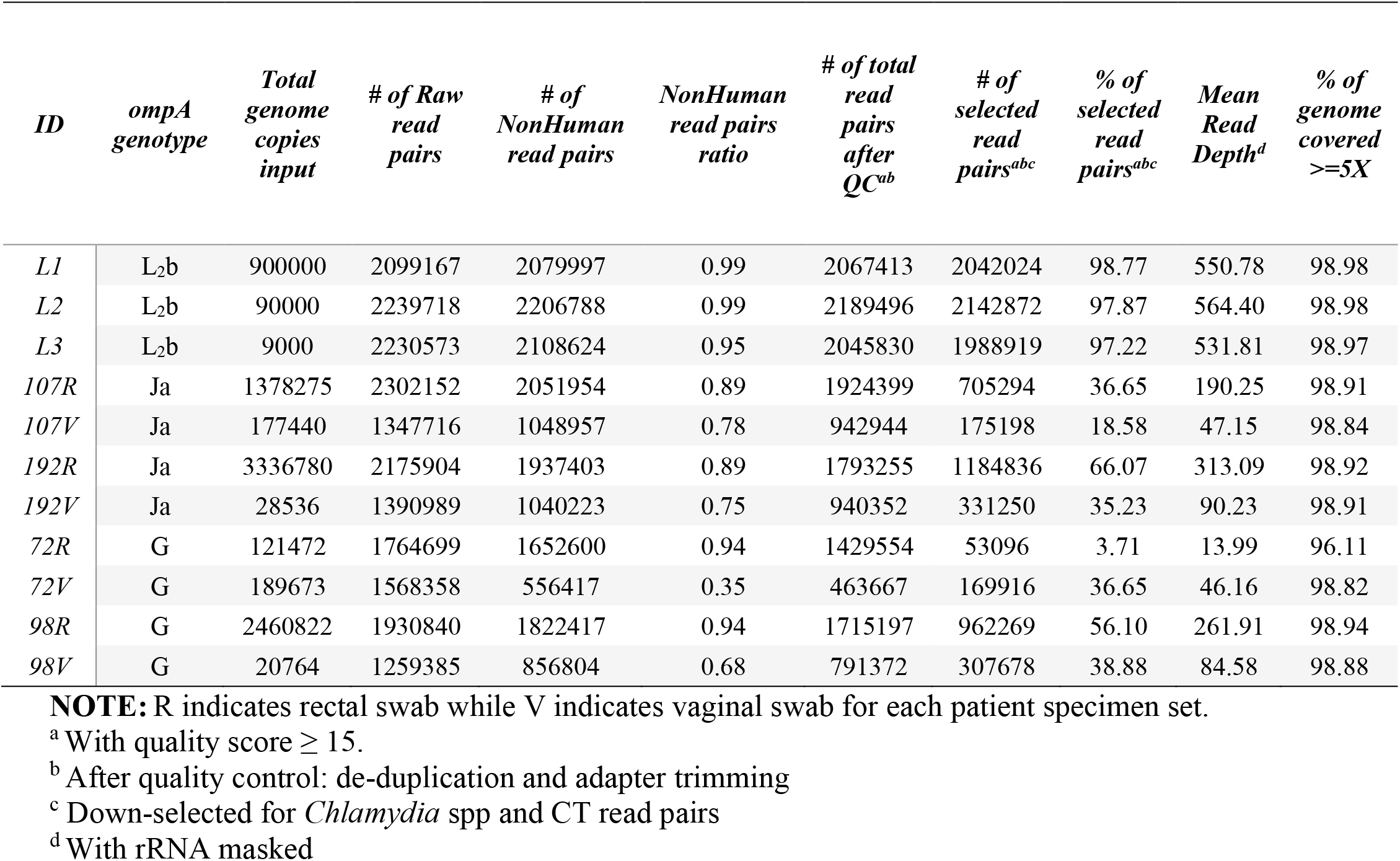
Sequence data analysis of *C. trachomatis* gDNA extracted from spiked mock samples of genotype L_2_b and patient specimen sets.

To determine efficiency of target enrichment from vaginal and rectal clinical specimens, *C. trachomatis* was directly sequenced from 4 sets of patient-matched vaginal and rectal swabs. The range in copy number by qPCR was 328 to 2,218 genomes per μL for the vaginal samples and 1,664 to 43,905 genomes per μL for the rectal samples. Total input ranged from 20,764 to 3,336,780 *C. trachomatis* genome copies. More genome copies were present in the rectal swab compared to the vaginal swabs in 3 of the 4 patient specimen sets (Table 2). As with the spiked gDNA serial dilutions, the proportion of reads classified as *Chlamydia* spp from clinical samples was dependent on total genome copy input (Table 2, Figure 2).

**Figure 2:**
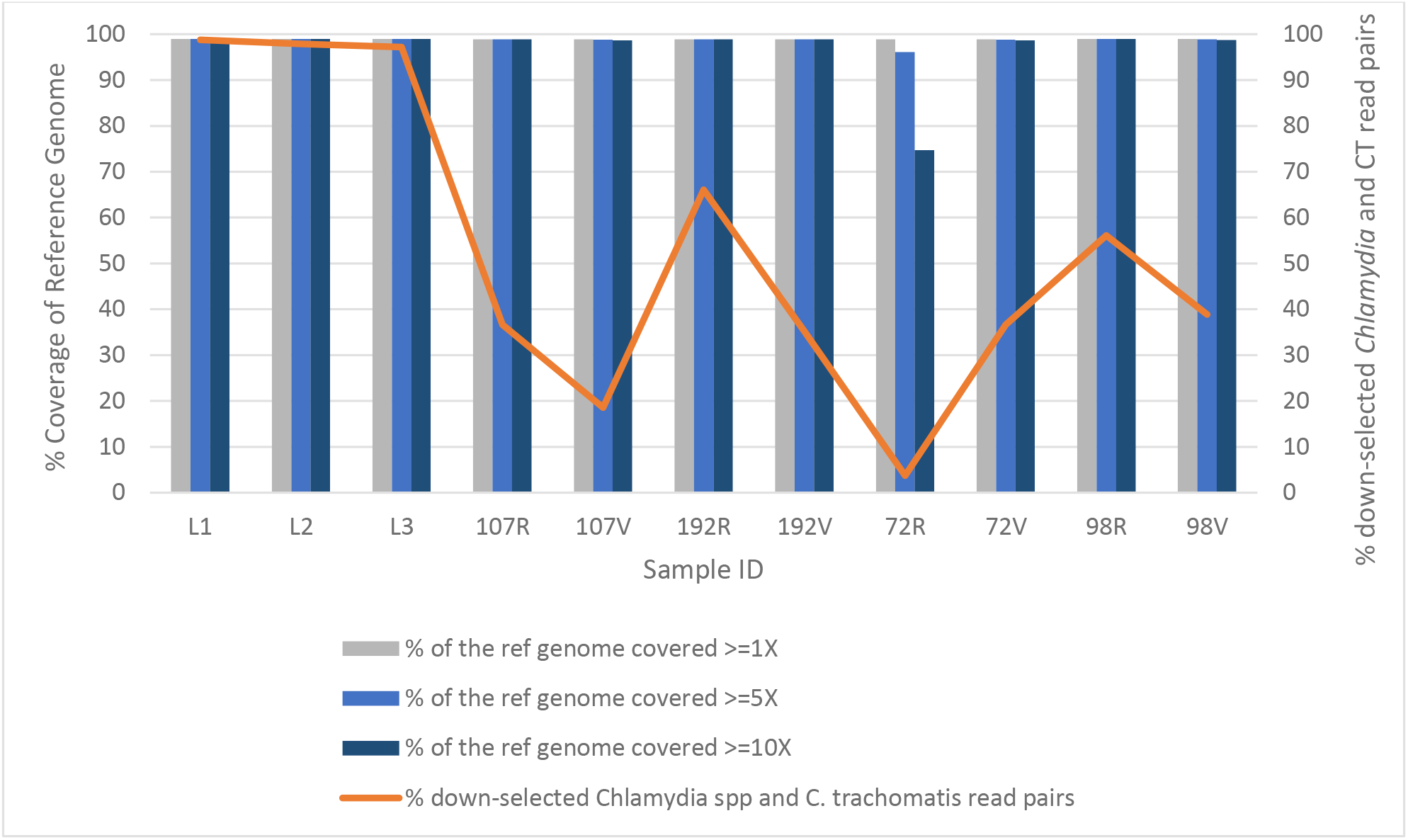
Percent coverage of reference genome and reads mapping to the respective reference genome for spiked mock samples of genotype L_2_b and patient specimen sets. The percent coverage of the reference genome for each patient specimen is represented as bars, with bar coloring based on average mean read depth as indicated. R indicates rectal sample while V indicates vaginal sample for each patient specimen set. The orange line represents the percent of down-selected *Chlamydia* spp and *C. trachomatis* read pairs.

For three of the four patient specimen sets 107, 192, and 98, 18.07%, 30.82%, and 17.13% more reads were classified, respectively, as *Chlamydia* spp in the rectal swab than the respective vaginal swab (Table 2, Figure 2). Interestingly, for patient specimen set 72, there were 32.94% more *Chlamydia* spp reads in the vaginal swab, which contained 68,201 more genome copies than the rectal swab (Table 2, Figure 2). Mean read depth was on average 3.5fold higher in rectal swabs from patient specimen sets 107, 192, and 98, while patient specimen set 72 demonstrated a 3.3fold higher mean read depth in the vaginal swab (Table 2; Supplementary Figure S2). For all patient specimen sets, the percentage of *C. trachomatis* genome covered with at least 5X coverage was > 98% with the exception of 72R, which had only 96.11% of the genome covered at a minimum of 5X read depth.

### Phylogeny and detection of recombination events from genomes derived from patient specimens

To ensure genomes generated from the four patient specimen sets clustered to any of the four deep-branching monophyletic *C. trachomatis* lineages, a whole genome phylogenetic analysis was constructed with 94 *C. trachomatis* single-contig genome sequences that were available from GenBank in December 2019 (Supplementary Table ST2)^40,42,55^. Both the genomes from patient specimen sets 107 and 192 clustered within the “prevalent urogenital and rectal strain” clade, while patient specimen sets 98 and 72 clustered within the “non-prevalent urogenital and rectal strain” clade (Figure 3).

**Figure 3:**
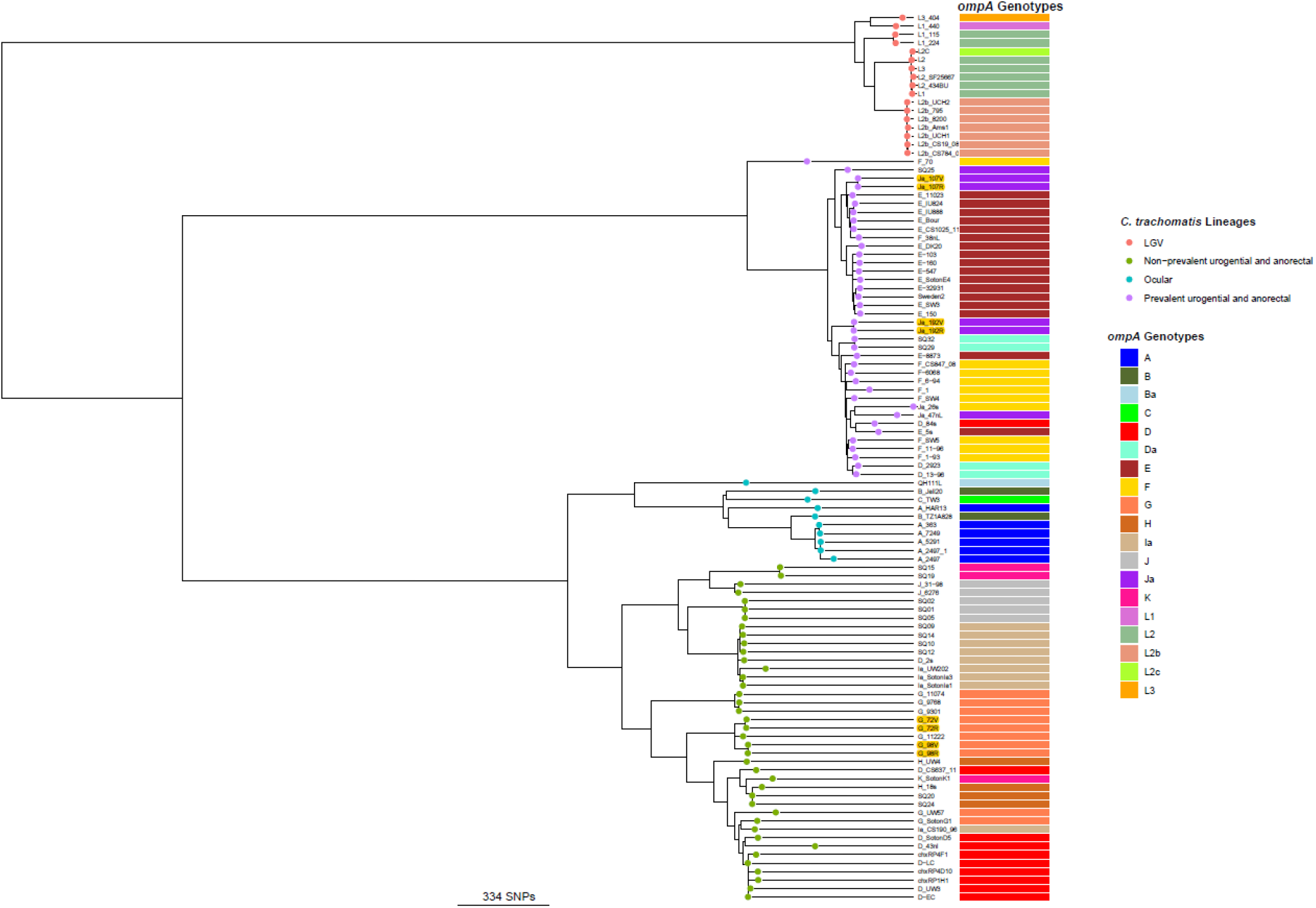
Global phylogeny of patient specimen sets and 94 *C. trachomatis* complete genomes. The four major lineages of *C. trachomatis* are highlighted with circular tip shapes in four distinct colors. The associated *ompA* genotypes for each of genome derived from the whole genomes data is also shown with the color code shown on the right-hand rectangular box. Patient specimen sets are highlighted in yellow.

The 107 and 192 patient genomes clustered in the same clade as the E genotype genomes (prevalent urogenital and anorectal lineage); whereas patient 98 and 72 genomes clustered in the same clade as G *ompA* genotypes (Non-prevalent urogenital and anorectal lineage) (Figure 3). For all four patients, the genomes derived from the two body sites formed distinct monophyletic clades within their respective urogenital lineage with a mean SNP difference of 3 SNPs (1-5 SNPs), indicating that each patient likely carried the same strain within their rectum and vagina (Figure 3, Table 3). Interestingly, five within host SNPs were identified in patient 72, and 2 of these SNPs were within the highly recombinogenic *ompA* (2 SNPs) and *pmpF* (2 SNPs) genes. Genomic comparison of all the plasmids against all reference plasmid sequences showed that within each patient specimen set there was 100% sequence similarity within the set (e.g., sample sets 107 and 192 had E plasmids in both anatomic sites) with the exception that the vaginal plasmid from patient 192 had a single nucleotide deletion at nucleotide position 5241 compared to the rectal plasmid. This deletion was within a gene that encodes a hypothetical protein. No other indels or SNPs were noted in any of the plasmids.

**Table 3:** Single nucleotide polymorphisms (SNPs) detected within patient specimen sets.

| *Patient specimen set* | *Genes containing within host SNPs* | *Number of SNPs* | *Genome location* | *Reference genome used for calling the SNPs<sup>c</sup>* |
| --- | --- | --- | --- | --- |
| 107R<sup>a</sup> and 107V<sup>b</sup> | Hypothetical protein (FSW4_RS00260) | 1 | 54906 | F/SW4 (NC_017951.1) |
|  | Hypothetical protein (FSW4_RS00270) | 1 | 59520, 59523 | F/SW4 (NC_017951.1) |
|  | Intergenic region | 1 | 79002 | F/SW4 (NC_017951.1) |
|  | Amino acid ABC transporter ATP-binding protein | 1 | 146249 | F/SW4 (NC_017951.1) |
|  | Intergenic region | 1 | 437237 | F/SW4 (NC_017951.1) |
| 192R and 192V | PTS sugar transporter subunit IIA | 1 | 326567 | F/SW4 (NC_017951.1) |
| 72R and 72V | *ompA* | 2 | 779230, 779235 | D/UW-3/CX (NC_000117.1) |
|  | Hypothetical protein (CT_744) | 1 | 864903 | D/UW-3/CX (NC_000117.1) |
|  | *pmpF* | 2 | 1029641, 1029643 | D/UW-3/CX (NC_000117.1) |
| 98R and 98V | Hypothetical protein (CT_049) | 1 | 54281 | D/UW-3/CX (NC_000117.1) |

<sup>a</sup>R, rectal <sup>b</sup>V, vaginal <sup>c</sup>SNPs were called by comparing the genomes of patient specimen sets to reference genomes that clustered in the whole genome phylogenetic tree.

Patient specimen sets harbored a total of 14 putative recombination blocks that contained between 21 and 419 homoplasic SNPs estimated to be introduced due to homologous recombination. The number of putative recombination blocks varied between 1 and 6 blocks within a patient specimen set, covering an average of 12.2 kb region per specimen (Supplementary Table ST3). All the putative recombination blocks identified and described were shared among and detected only in each of the patient specimen sets indicating that these recombination events were ancestral and acquired through clonal descent. Among the patient specimen sets, 107R and 107V (urogenital prevalent lineage) had the highest rates of recombination (ρ/θ = 0.111) as well as increased effects of recombination over point mutations (r/m = 5.777) followed by patient specimen set 72R and 72V (urogenital non-prevalent lineage; ρ/θ = 0.053; r/m = 3.803). The lowest number of recombination events were observed among the 98V-98R patient specimen sets (non-prevalent lineage; ρ/θ = 0.031; r/m = 0.6562) (Supplementary Table ST3).

Some of the previously identified genomic regions of higher homologous recombination were also identified in this study. The *ompA* gene has undergone recombination in both of the prevalent urogenital 107 and 192 patient specimen sets. *ompA* genotyping by both Sanger sequencing and whole genome data indicated the *ompA* genotype of specimens derived from patients 107 and 192 was genotype Ja within an E genome backbone. Our recombination detection analysis showed that homologous recombination might have mediated the transfer of a ~12.9Kbp fragment containing CT_681/*ompA* along with the neighboring genes (from Type III secretion system protein gene [CT_672] to *pbpB* [CT_682]) into the 192 patient and a larger fragment of 17.6Kbp from CT_672 to *pbpB* (CT_682) along with *ompA* into the 107 patient, likely from a Ja strain (Figure 3; Supplementary Table ST3). The *pmpE* and *mrsA_1* genes were estimated to be recombinant, respectively, in sets 107 and 192^40,55^. The gene encoding the inclusion membrane protein gene *incD* was only predicted to be recombinant in the non-prevalent urogenital patient specimen set 98. (Supplementary Table ST3)^40,55^.

## Discussion

A SureSelect^XT^ workflow with an expanded 2.698 Mbp RNA bait library with 34,795, 120-mer probes was developed from 85 GenBank *C. trachomatis* reference genomes encompassing all four lineages of *C. trachomatis* compared to the previous RNA bait library developed from 74 GenBank *C. trachomatis* reference genomes with 33,619, 120-mer probes^52^. Our expanded library was used to enrich and sequence eight *C. trachomatis* genomes directly from paired clinical vaginal and rectal swabs from the same four patients, respectively. Although the RNA bait library developed here did not contain plasmid sequences as an oversight, compared to a previous study that obtained 98% coverage with a total input of 4,800 *C. trachomatis* strain F/SW4 genome copies^52^, we report an LOD of around 16,000 total genomes at 98% coverage for *C. trachomatis* reference strain D/UW-3/Cx. This LOD was within the same order of magnitude as the previous study, although we utilized an expanded RNA bait library, a different assay to determine copy number (i.e., qRT-PCR versus qPCR), and gDNA from a spiked sample versus a sample that had been propagated in tissue culture.

In this study, it was useful to calculate comparable mean read depths and the number of down-selected read pairs among the spiked samples of L_2_b, as this demonstrates success in enrichment for LGV strains as they have become prevalent among MSM^23,25,27,28^. Phylogenetic analysis of the 94 reference genomes (of which 85 genomes were used to develop the RNA bait library) showed genome representation from all four *C. trachomatis* lineages (Figure 3). Overall, this workflow was successful in enrichment and sequencing of *C. trachomatis* strains that are prevalent in anorectal infections from two different populations: the MSM population from which L_2_b originated and the female heterosexual population from which the paired vaginal and rectal samples originated.

The sequence data analysis of patient specimen sets of vaginal and rectal swabs revealed successful enrichment and genome sequencing of *C. trachomatis* from both clinical specimen types. In three of the four patient specimen sets, sequencing was more efficient for the rectal swabs, likely due to the higher genome copy number of *C. trachomatis* for those specimen sets. With a total input of 3,336,780 *C. trachomatis* genome copies, the rectal patient specimen 192 demonstrated the best enrichment with 89% nonhuman reads, of which 66.07% belonged to *Chlamydia* species that mapped to 98.92% of the *C. trachomatis* reference E strain genome with a mean read depth coverage of 313.09. This is an improvement over a previous study that used clinical urine and vaginal samples where the best sample had 49.57% of the reads belonging to *Chlamydia* species with 99.9% mapping to the reference D genome at a 410 mean read depth from a total input of 68,864,400 *C. trachomatis* genome copies^52^. Overall, enrichment of *C. trachomatis* from the samples in that study was successful in only eight (80%) of 10 samples for a genome coverage of ≥95-100% for the respective reference genome; the percentage of reads mapping to *C. trachomatis* ranged from 0.07-49.57% across the specimens possibly due to hybridization of the RNA bait library primers with human DNA. In another study using a similar bait library, only 12 (60%) of 20 clinical ocular samples reached a ≥95-100% genome coverage^56^. In our study, all eight (100%) clinical samples reached a ≥95-100% genome coverage with reads mapping to *C. trachomatis* ranging from 3.71-98.77%, indicating that the expanded probe library—that further excludes baits with human homology—can achieve the desired efficiency.

The higher *C. trachomatis* bacterial load detected in rectal specimens in three of the four patient specimen sets conflicts with two studies that demonstrated similar loads across sets of vaginal and anorectal specimens collected from the same women with and without anal intercourse visiting a STI clinic in the Netherlands and in a high HIV prevalence area in South Africa^76,77^. For this specific study, the trend may be a characteristic of the assay used to determine copy number, the study population itself or, because of the small sample size, may not be representative of the study population as a whole. Nevertheless, Dirks et al.^78^ pointed out the ineffectiveness of comparing load across *C. trachomatis* surveillance studies due to the lack of standardization for load determination and the presence of inflammatory cells which can artificially lower the number of *C. trachomatis* load. However, it is useful to determine the bacterial load in patient specimens to ensure enough gDNA is present to successfully enrich for *C. trachomatis* in a target-enrichment sequencing workflow, although conclusions should not be drawn from this estimated value about severity of infection, associated symptoms, or transmission.

The genomes derived from each patient specimen set were phylogenetically associated within the representative lineages of their respective strain as determined by Sanger sequencing, demonstrating success of the workflow and bioinformatic analysis described here. Interestingly, patient specimen sets 107 and 192 were associated with the prevalent lineage, a recent urogenital lineage that has been suggested to have been derived from recombination^40,55^. Further, the detection of recombinant *ompA,* a known hotspot for recombination in the *C. trachomatis* genome, in these samples as part of an approximate 12kb exchange could have only been identified by WGS and not by Sanger sequencing alone, highlighting the genetic resolution achieved in this study to understand the evolution of contemporary *C. trachomatis* strains^40,43,55^. Some studies have reported recombinant forms of both *ompA* and *pmpH,* and tracking these and additional known recombinant variants may inform the field on the dynamics of current transmission networks and *C. trachomatis* tropism in general^9,53,79^. With that said, the high rates of recombination and limited genetic diversity within *C. trachomatis* strains indicates the need to include a large number of loci dispersed throughout the genome to circumvent false negativity of clinical strain typing, which can now be efficiently achieved by sequencing the whole genome.

The circulation of two different *C. trachomatis* lineages with varying degrees of recombination across the genomes in a single population is of interest and displays the complexity of *C. trachomatis* strain evolution and transmission, which cannot be discerned with traditional molecular typing methods such as *ompA* genotyping, MLST or MLVA. Here, we have described a higher efficiency target-enrichment bait library that streamlines the molecular characterization of *C. trachomatis* from rectal as well as vaginal specimens. This type of high-resolution data can be used to understand the genetic diversity of current *C. trachomatis* strains causing genital and rectal infections and provide a robust molecular epidemiologic approach to advance our understanding of geo-sexual networks, outbreaks of colorectal infections among women and men who have sex with men, and the role of these strains in morbidity. The bioinformatic pipeline can further be used to potentially identify novel markers for typing *C. trachomatis* and to examine the microbiome to determine the role it plays in susceptibility, transmission and clearance of rectal *C. trachomatis* infections, especially given the need for a longer duration of therapy compared to most uncomplicated urogenital infections^80,81^.

## Acknowledgements

The findings and conclusions in this report are those of the authors and do not necessarily represent the official position of the Centers for Disease Control and Prevention. This work was made possible through support from CDC’s Advanced Molecular Detection (AMD) program. The funders had no role in study design, data collection and interpretation, or the decision to submit the work for publication.

We thank Sankhya Bomanna, PhD, for excellent technical assistance.

## Authors’ information

K. Bowden is now at the Division of Parasitic Diseases and Malaria, Centers for Disease Control and Prevention, Atlanta, GA, USA

## Author Contribution Statement

K. Bowden transcribed the manuscript and generated the genomic libraries from the serial dilutions and clinical specimens for sequencing and analysis.

S. Joseph performed the bioinformatics analysis, conducted phylogenetic analysis of the genomic data and contributed to the writing of the manuscript.

J. Cartee pooled and sequenced the genomic libraries on the MiSeq.

N. Ziklo prepared and purified the gDNA from the clinical samples and performed the qPCR and fragmentation for library preparation.

D. Danavall prepared spiked gDNA serial dilutions for LOD determination.

B. Raphael oversaw the project within CDC and provided technical support and subject matter expertise on the genome sequencing workflow and bioinformatic analysis.

T. Read conducted initial phylogenetic analysis while providing continued technical support and subject matter expertise for the data analysis.

D. Dean collected and provided the patient specimens and provided technical support and subject matter expertise on the genome sequencing workflow and contributed to the writing of the manuscript.

All authors reviewed, edited, and contributed to the manuscript.

## Additional Information

The authors declare no competing interests.

## Supplementary Information

ST1. Primers used for PCR amplification and sequencing of the entire *C. trachomatis* plasmid for each patient sample set.

ST2. List of *C. trachomatis* genomes and their NCBI/SRA accession IDs used for reconstructing the global phylogenetic tree and subsequent putative recombination detection analysis.

ST3. Genes present in putative recombination blocks for patient specimen sets.

S1. Coverage plots for reference strain D/UW-3/Cx.

S2. Coverage plots for L_2_b strain and patient specimen sets.

## References

1. Centers for Disease C, Prevention. Recommendations for the laboratory-based detection of *Chlamydia trachomatis* and *Neisseria gonorrhoeae*--2014. MMWR Recomm Rep. 2014;63(RR-02):1–19.

2. Stamm WE. *Chlamydia trachomatis* infections of the adult. In: Holmes KK, Sparling FF, Stamm WE., et al., eds. Sexually transmitted diseases. Vol 4th. 4th ed. New York: McGraw Hill Professional; 2011: http://catdir.loc.gov/catdir/toc/ecip0714/2007013008.html

3. Dean D, Schachter J, Dawson CR, Stephens RS. Comparison of the major outer membrane protein variant sequence regions of B/Ba isolates: a molecular epidemiologic approach to *Chlamydia trachomatis* infections. J Infect Dis. 1992;166(2):383–392.

4. Isaksson J, Carlsson O, Airell A, Stromdahl S, Bratt G, Herrmann B. Lymphogranuloma venereum rates increased and *Chlamydia trachomatis* genotypes changed among men who have sex with men in Sweden 2004-2016. J Med Microbiol. 2017;66(11):1684–1687.

5. Spaargaren J, Fennema HS, Morre SA, de Vries HJ, Coutinho RA. New lymphogranuloma venereum *Chlamydia trachomatis* variant, Amsterdam. Emerg Infect Dis. 2005;11(7):1090–1092.

6. Spaargaren J, Schachter J, Moncada J, et al. Slow epidemic of lymphogranuloma venereum L2b strain. Emerg Infect Dis. 2005;11(11):1787–1788.

7. Christerson L, de Vries HJ, de Barbeyrac B, et al. Typing of lymphogranuloma venereum *Chlamydia trachomatis* strains. Emerg Infect Dis. 2010;16(11):1777–1779.

8. Versteeg B, van Rooijen MS, Schim van der Loeff MF, de Vries HJC, Bruisten SM. No indication for tissue tropism in urogenital and anorectal *Chlamydia trachomatis* infections using high-resolution multilocus sequence typing. BMC Infectious Diseases. 2014;14(1):464.

9. Bom RJ, van der Helm JJ, Schim van der Loeff MF, et al. Distinct transmission networks of *Chlamydia trachomatis* in men who have sex with men and heterosexual adults in Amsterdam, The Netherlands. PLoS One. 2013;8(1):e53869.

10. Lysen M, Osterlund A, Rubin CJ, Persson T, Persson I, Herrmann B. Characterization of *ompA* genotypes by sequence analysis of DNA from all detected cases of *Chlamydia trachomatis* infections during 1 year of contact tracing in a Swedish County. J Clin Microbiol. 2004;42(4):1641–1647.

11. Quint KD, Bom RJ, Quint WG, et al. Anal infections with concomitant *Chlamydia trachomatis* genotypes among men who have sex with men in Amsterdam, the Netherlands. BMC Infect Dis. 2011;11:63.

12. Bax CJ, Quint KD, Peters RP, et al. Analyses of multiple-site and concurrent *Chlamydia trachomatis* serovar infections, and serovar tissue tropism for urogenital versus rectal specimens in male and female patients. Sex Transm Infect. 2011;87(6):503–507.

13. Dewart CM, Bernstein KT, DeGroote NP, Romaguera R, Turner AN. Prevalence of Rectal Chlamydial and Gonococcal Infections: A Systematic Review. Sex Transm Dis. 2018;45(5):287–293.

14. Andersson N, Boman J, Nylander E. Rectal chlamydia - should screening be recommended in women? International journal of STD & AIDS. 2017;28(5):476–479.

15. Waalboer R, van der Snoek EM, van der Meijden WI, Mulder PG, Ossewaarde JM. Analysis of rectal *Chlamydia trachomatis* serovar distribution including L2 (lymphogranuloma venereum) at the Erasmus MC STI clinic, Rotterdam. Sex Transm Infect. 2006;82(3):207–211.

16. Morre SA, Rozendaal L, van Valkengoed IG, et al. Urogenital *Chlamydia trachomatis* serovars in men and women with a symptomatic or asymptomatic infection: an association with clinical manifestations? J Clin Microbiol. 2000;38(6):2292–2296.

17. Klint M, Lofdahl M, Ek C, Airell A, Berglund T, Herrmann B. Lymphogranuloma venereum prevalence in Sweden among men who have sex with men and characterization of *Chlamydia trachomatis ompA* genotypes. J Clin Microbiol. 2006;44(11):4066–4071.

18. Koper NE, van der Sande MA, Gotz HM, Koedijk FD. Lymphogranuloma venereum among men who have sex with men in the Netherlands: regional differences in testing rates lead to underestimation of the incidence, 2006-2012. Euro surveillance: bulletin Europeen sur les maladies transmissibles = European communicable disease bulletin. 2013;18(34).

19. Ward H, Alexander S, Carder C, et al. The prevalence of lymphogranuloma venereum infection in men who have sex with men: results of a multicentre case finding study. Sex Transm Infect. 2009;85(3):173–175.

20. Stark D, van Hal S, Hillman R, Harkness J, Marriott D. Lymphogranuloma venereum in Australia: anorectal *Chlamydia trachomatis* serovar L2b in men who have sex with men. J Clin Microbiol. 2007;45(3):1029–1031.

21. Ronn MM, Ward H. The association between lymphogranuloma venereum and HIV among men who have sex with men: systematic review and meta-analysis. BMC Infect Dis. 2011;11:70.

22. Chan PA, Robinette A, Montgomery M, et al. Extragenital Infections Caused by *Chlamydia trachomatis* and *Neisseria gonorrhoeae*: A Review of the Literature. Infectious diseases in obstetrics and gynecology. 2016;2016:5758387.

23. van Liere GA, van Rooijen MS, Hoebe CJ, Heijman T, de Vries HJ, Dukers-Muijrers NH. Prevalence of and Factors Associated with Rectal-Only Chlamydia and Gonorrhoea in Women and in Men Who Have Sex with Men. PLoS One. 2015;10(10):e0140297.

24. Chandra NL, Broad C, Folkard K, et al. Detection of *Chlamydia trachomatis* in rectal specimens in women and its association with anal intercourse: a systematic review and meta-analysis. Sex Transm Infect. 2018;94(5):320–326.

25. Danby CS, Cosentino LA, Rabe LK, et al. Patterns of Extragenital Chlamydia and Gonorrhea in Women and Men Who Have Sex With Men Reporting a History of Receptive Anal Intercourse. Sex Transm Dis. 2016;43(2):105–109.

26. Kong FY, Tabrizi SN, Law M, et al. Azithromycin versus doxycycline for the treatment of genital chlamydia infection: a meta-analysis of randomized controlled trials. Clin Infect Dis. 2014;59(2):193–205.

27. Foschi C, Gaspari V, Sgubbi P, Salvo M, D’Antuono A, Marangoni A. Sexually transmitted rectal infections in a cohort of ‘men having sex with men’. J Med Microbiol. 2018;67(8):1050–1057.

28. de Vrieze NH, de Vries HJ. Lymphogranuloma venereum among men who have sex with men. An epidemiological and clinical review. Expert Rev Anti Infect Ther. 2014;12(6):697–704.

29. Bachmann LH, Johnson RE, Cheng H, et al. Nucleic acid amplification tests for diagnosis of *Neisseria gonorrhoeae* and *Chlamydia trachomatis* rectal infections. J Clin Microbiol. 2010;48(5):1827–1832.

30. Labiran C, Marsh P, Zhou J, et al. Highly diverse MLVA-ompA genotypes of rectal *Chlamydia trachomatis* among men who have sex with men in Brighton, UK and evidence for an HIV-related sexual network. Sex Transm Infect. 2016;92(4):299–304.

31. Labiran C, Rowen D, Clarke IN, Marsh P. Detailed molecular epidemiology of *Chlamydia trachomatis* in the population of Southampton attending the genitourinary medicine clinic in 2012-13 reveals the presence of long established genotypes and transitory sexual networks. PLoS One. 2017;12(9):e0185059.

32. Dean D, Bruno WJ, Wan R, et al. Predicting phenotype and emerging strains among *Chlamydia trachomatis* infections. Emerg Infect Dis. 2009;15(9):1385–1394.

33. Kapil R, Press CG, Hwang ML, Brown L, Geisler WM. Investigating the epidemiology of repeat *Chlamydia trachomatis* detection after treatment by using C. trachomatis OmpA genotyping. J Clin Microbiol. 2015;53(2):546–549.

34. Herrmann B, Isaksson J, Ryberg M, et al. Global Multilocus Sequence Type Analysis of *Chlamydia trachomatis* Strains from 16 Countries. J Clin Microbiol. 2015;53(7):2172–2179.

35. Pannekoek Y, Morelli G, Kusecek B, et al. Multi locus sequence typing of Chlamydiales: clonal groupings within the obligate intracellular bacteria *Chlamydia trachomatis*. BMC Microbiol. 2008;8:42.

36. Bom RJ, Christerson L, Schim van der Loeff MF, Coutinho RA, Herrmann B, Bruisten SM. Evaluation of high-resolution typing methods for *Chlamydia trachomatis* in samples from heterosexual couples. J Clin Microbiol. 2011;49(8):2844–2853.

37. Klint M, Fuxelius HH, Goldkuhl RR, et al. High-resolution genotyping of *Chlamydia trachomatis* strains by multilocus sequence analysis. J Clin Microbiol. 2007;45(5):1410–1414.

38. Lan J, Walboomers JM, Roosendaal R, et al. Direct detection and genotyping of *Chlamydia trachomatis* in cervical scrapes by using polymerase chain reaction and restriction fragment length polymorphism analysis. J Clin Microbiol. 1993;31(5):1060–1065.

39. Smelov V, Vrbanac A, van Ess EF, et al. *Chlamydia trachomatis* Strain Types Have Diversified Regionally and Globally with Evidence for Recombination across Geographic Divides. Front Microbiol. 2017;8:2195.

40. Harris SR, Clarke IN, Seth-Smith HM, et al. Whole-genome analysis of diverse *Chlamydia trachomatis* strains identifies phylogenetic relationships masked by current clinical typing. Nat Genet. 2012;44(4):413–419, S411.

41. Joseph SJ, Didelot X, Gandhi K, Dean D, Read TD. Interplay of recombination and selection in the genomes of *Chlamydia trachomatis*. Biol Direct. 2011;6:28.

42. Joseph SJ, Read TD. Genome-wide recombination in *Chlamydia trachomatis*. Nat Genet. 2012;44(4):364–366.

43. Gomes JP, Bruno WJ, Nunes A, et al. Evolution of *Chlamydia trachomatis* diversity occurs by widespread interstrain recombination involving hotspots. Genome Res. 2007;17(1):50–60.

44. Somboonna N, Wan R, Ojcius DM, et al. Hypervirulent *Chlamydia trachomatis* clinical strain is a recombinant between lymphogranuloma venereum (L(2)) and D lineages. mBio. 2011;2(3):e00045–00011.

45. Castro R, Baptista T, Vale A, et al. Lymphogranuloma venereum serovar L2b in Portugal. International journal of STD & AIDS. 2010;21(4):265–266.

46. Martin-Iguacel R, Llibre JM, Nielsen H, et al. Lymphogranuloma venereum proctocolitis: a silent endemic disease in men who have sex with men in industrialised countries. Eur J Clin Microbiol Infect Dis. 2010;29(8):917–925.

47. Borges V, Cordeiro D, Salas AI, et al. *Chlamydia trachomatis*: when the virulence-associated genome backbone imports a prevalence-associated major antigen signature. Microbial genomics. 2019;5(11).

48. Taylor-Brown A, Madden D, Polkinghorne A. Culture-independent approaches to chlamydial genomics. Microbial genomics. 2018;4(2).

49. Putman TE, Suchland RJ, Ivanovitch JD, Rockey DD. Culture-independent sequence analysis of *Chlamydia trachomatis* in urogenital specimens identifies regions of recombination and in-patient sequence mutations. Microbiology. 2013;159(Pt 10):2109–2117.

50. Hedrum A, Lundeberg J, Pahlson C, Uhlen M. Immunomagnetic recovery of *Chlamydia trachomatis* from urine with subsequent colorimetric DNA detection. PCR Methods Appl. 1992;2(2):167–171.

51. Seth-Smith HM, Harris SR, Skilton RJ, et al. Whole-genome sequences of *Chlamydia trachomatis* directly from clinical samples without culture. Genome Res. 2013;23(5):855–866.

52. Christiansen MT, Brown AC, Kundu S, et al. Whole-genome enrichment and sequencing of *Chlamydia trachomatis* directly from clinical samples. BMC Infect Dis. 2014;14:591.

53. Borges V, Cordeiro D, Salas AI, et al. *Chlamydia trachomatis* outbreak: when the virulence-associated genome backbone imports a prevalence-associated major antigen signature. 2019:622324.

54. Last AR, Pickering H, Roberts CH, et al. Population-based analysis of ocular *Chlamydia trachomatis* in trachoma-endemic West African communities identifies genomic markers of disease severity. Genome Med. 2018;10(1):15.

55. Hadfield J, Harris SR, Seth-Smith HMB, et al. Comprehensive global genome dynamics of *Chlamydia trachomatis* show ancient diversification followed by contemporary mixing and recent lineage expansion. Genome Res. 2017;27(7):1220–1229.

56. Alkhidir AAI, Holland MJ, Elhag WI, et al. Whole-genome sequencing of ocular *Chlamydia trachomatis* isolates from t Vectors. 2019;12(1):518.

57. Pickering H, Chernet A, Sata E, et al. Genomics of Ocular *Chlamydia trachomatis* after 5 years of SAFE interventions for trachoma in Amhara, Ethiopia. 2020:2020.2006.2007.138982.

58. Chen CY, Chi KH, Alexander S, Ison CA, Ballard RC. A real-time quadriplex PCR assay for the diagnosis of rectal lymphogranuloma venereum and non-lymphogranuloma venereum *Chlamydia trachomatis* infections. Sex Transm Infect. 2008;84(4):273–276.

59. Ravel J, Gajer P, Abdo Z, et al. Vaginal microbiome of reproductive-age women. Proc Natl Acad Sci U S A. 2011;108 Suppl 1:4680–4687.

60. Sharma M, Recuero-Checa MA, Fan FY, Dean D. *Chlamydia trachomatis* regulates growth and development in response to host cell fatty acid availability in the absence of lipid droplets. Cell Microbiol. 2018;20(2).

61. Batteiger BE, Wan R, Williams JA, et al. Novel *Chlamydia trachomatis* strains in heterosexual sex partners, Indianapolis, Indiana, USA. Emerg Infect Dis. 2014;20(11):1841–1847.

62. Katoh K, Standley DM. MAFFT multiple sequence alignment software version 7: improvements in performance and usability. Mol Biol Evol. 2013;30(4):772–780.

63. Langmead B, Salzberg SL. Fast gapped-read alignment with Bowtie 2. Nat Methods. 2012;9(4):357–359.

64. Sachidanandam R, Weissman D, Schmidt SC, et al. A map of human genome sequence variation containing 1.42 million single nucleotide polymorphisms. Nature. 2001;409(6822):928–933.

65. Martin M. Cutadapt removes adapter sequences from high-throughput sequencing reads. 2011. 2011;17(1):3 J EMBnet.journal.

66. Ainsworth D, Sternberg MJE, Raczy C, Butcher SA. k-SLAM: accurate and ultra-fast taxonomic classification and gene identification for large metagenomic data sets. Nucleic Acids Res. 2017;45(4):1649–1656.

67. Truong DT, Franzosa EA, Tickle TL, et al. MetaPhlAn2 for enhanced metagenomic taxonomic profiling. Nat Methods. 2015;12(10):902–903.

68. Quinlan AR, Hall IM. BEDTools: a flexible suite of utilities for comparing genomic features. Bioinformatics. 2010;26(6):841–842.

69. Li H. Aligning sequence reads, clone sequences and assembly contigs with BWA-MEM. arXiv e-prints. 2013. https://ui.adsabs.harvard.edu/abs/2013arXiv1303.3997L. Accessed March 01, 2013.

70. Li H, Handsaker B, Wysoker A, et al. The Sequence Alignment/Map format and SAMtools. Bioinformatics. 2009;25(16):2078–2079.

71. Bankevich A, Nurk S, Antipov D, et al. SPAdes: a new genome assembly algorithm and its applications to single-cell sequencing. J Comput Biol. 2012;19(5):455–477.

72. Altschul SF, Gish W, Miller W, Myers EW, Lipman DJ. Basic local alignment search tool. J Mol Biol. 1990;215(3):403–410.

73. Garrison E, Marth G. Haplotype-based variant detection from short-read sequencing. arXiv e-prints. 2012. https://ui.adsabs.harvard.edu/abs/2012arXiv1207.3907G. Accessed July 01, 2012.

74. Croucher NJ, Page AJ, Connor TR, et al. Rapid phylogenetic analysis of large samples of recombinant bacterial whole genome sequences using Gubbins. Nucleic Acids Res. 2015;43(3):e15.

75. Stamatakis A. RAxML version 8: a tool for phylogenetic analysis and post-analysis of large phylogenies. Bioinformatics. 2014;30(9):1312–1313.

76. Dirks JAMC, van Liere GAFS, Hoebe CJPA, Wolffs P, Dukers-Muijrers NHTM. Genital and anal *Chlamydia trachomatis* bacterial load in concurrently infected women: a cross-sectional study. 2019;95(5):317–321.

77. Dubbink JH, de Waaij DJ, Bos M, et al. Microbiological Characteristics of *Chlamydia trachomatis* and *Neisseria gonorrhoeae* Infections in South African Women. Journal of clinical microbiology. 2016;54(1):200–203.

78. Dirks JAMC, Hoebe CJPA, van Liere GAFS, Dukers-Muijrers NHTM, Wolffs PFG. Standardisation is necessary in urogenital and extragenital *Chlamydia trachomatis* bacterial load determination by quantitative PCR: a review of literature and retrospective study. 2019:sextrans-2018-053522.

79. Rodriguez-Dominguez M, Gonzalez-Alba JM, Puerta T, et al. Spread of a new *Chlamydia trachomatis* variant from men who have sex with men to the heterosexual population after replacement and recombination in ompA and pmpH genes. Clinical microbiology and infection: the official publication of the European Society of Clinical Microbiology and Infectious Diseases. 2017;23(10):761–766.

80. Dukers-Muijrers NH, Speksnijder AG, Morre SA, et al. Detection of anorectal and cervicovaginal *Chlamydia trachomatis* infections following azithromycin treatment: prospective cohort study with multiple time-sequential measures of rRNA, DNA, quantitative load and symptoms. PLoS One. 2013;8(11):e81236.

81. Kong FY, Hocking JS. Treatment challenges for urogenital and anorectal *Chlamydia trachomatis*. BMC Infect Dis. 2015;15:293.

